# Push-to-open: The Gating Mechanism of the Tethered Mechanosensitive Ion Channel NompC

**DOI:** 10.1101/853721

**Authors:** Yang Wang, Yifeng Guo, Chunhong Liu, Lei Wang, Aihua Zhang, Zhiqiang Yan, Chen Song

**Author notes:** These authors contributed equally to this work (Yang Wang, Yifeng Guo). Correspondence and requests for materials should be addressed to (CS), (ZY).

## Abstract

NompC was one of the earliest identified mechanosensitive ion channels responsible for the sensation of touch and balance in *Drosophila melanogaster*. A tethered gating model was proposed for NompC and the Cryo-EM structure has been solved. However, the atomistic mechano-gating mechanism still remains elusive. Here we show the atomistic details of the NompC channel opening in response to the compression of the intracellular domain while remaining closed under an intracellular stretch. This is demonstrated by all-atom molecular dynamics simulations and evidenced by electrophysiological experiments. Under intracellular compression, the ankyrin repeat region undergoes a significant conformational change and passes the mechanical force to the linker helices like a spring with a force constant of ~3.3 pN/nm. The linker helix region acts as a bridge between the ankyrin repeats and TRP domain, and most of the mutations breaking the hydrogen bonds around this region lead to the loss-of-function of the channel. Eventually, the compression-induced mechanical force is passed from the linker helices onto the TRP domain, which then undergoes a clockwise rotation that leads to the opening of the channel. This work provides a clear picture of how a pushing force opens the mechanosensitive ion channel NompC, which might be a universal gating mechanism of similar tethered mechanosensitive ion channels, enabling cells to feel and respond to compression or shrinking.

## Main Text

Many types of sensations initiate from the gating of transient receptor potential (TRP) ion channels, which regulates the intracellular cation concentration that triggers downstream signals (*1*–*6*). NompC is one of the earliest identified mechanosensitive ion channels belonging to the TRP family, which plays crucial roles in the sensation of light touch, hearing, balance, and locomotion of *Drosophila melanogaster* (*7*–*11*). NompC is structurally unique having the largest number of ankyrin repeats (ARs) among the known TRP channels (*3*), 29 in total. The AR region was found to be associated with microtubules and was proposed to act as a gating spring to regulate the channel states: the so-called “tethered gating model” (*12*–*14*). Although NompC orthologs have not been found in mammal species (*15*, *16*), it was shown to function in mechanosensation of *C. elegans and D. rerio* as well (*17*), and therefore, it serves as an ideal model for studying the molecular mechanism of the tethered mechano-gating. Recently, the Cryo-EM structure of NompC was resolved (*18*), which showed that four AR chains form a ~15 nm-long supercoiled helix and connect to the transmembrane (TM) pore domain via a linker helix (LH) region (Fig. 1A). Thus, the new structure further confirmed that the AR helices probably act like a spring to conduct forces to the TM pore when the neuron cells deform. However, what kind of forces (or what type of cell deformation) can open the NompC channel, and how the force is transduced from ARs to the transmembrane region to finally open the pore, were still elusive. In the previous studies, it was often implied that pulling the AR spring may open the channel (*14*, *19*). In contrast, there were also models indicating that a pushing force may be required to open the channel (*20*). Notably, a recent continuum mechanics study showed that the AR region might exert a clockwise torque on the linker helix under compression (looking from the intracellular side), and this could lead to a clockwise motion of the TRP domain that can potentially open the channel (*21*). Nonetheless, the detailed gating mechanism of this unique tethered ion channel remains unclear to date. Additionally, the membrane-surface-tension-induced ion channel gating provided a mechanism by which cells can respond to volume expansion (*22*–*26*), but there is no known mechano-gating mechanism that is able to respond to cell compression or volume shrinking. This work provides a plausible push-to-open mechanism for the tethered ion channels, which may be used by cells as a response to compression and shrinking.

**Figure 1.**
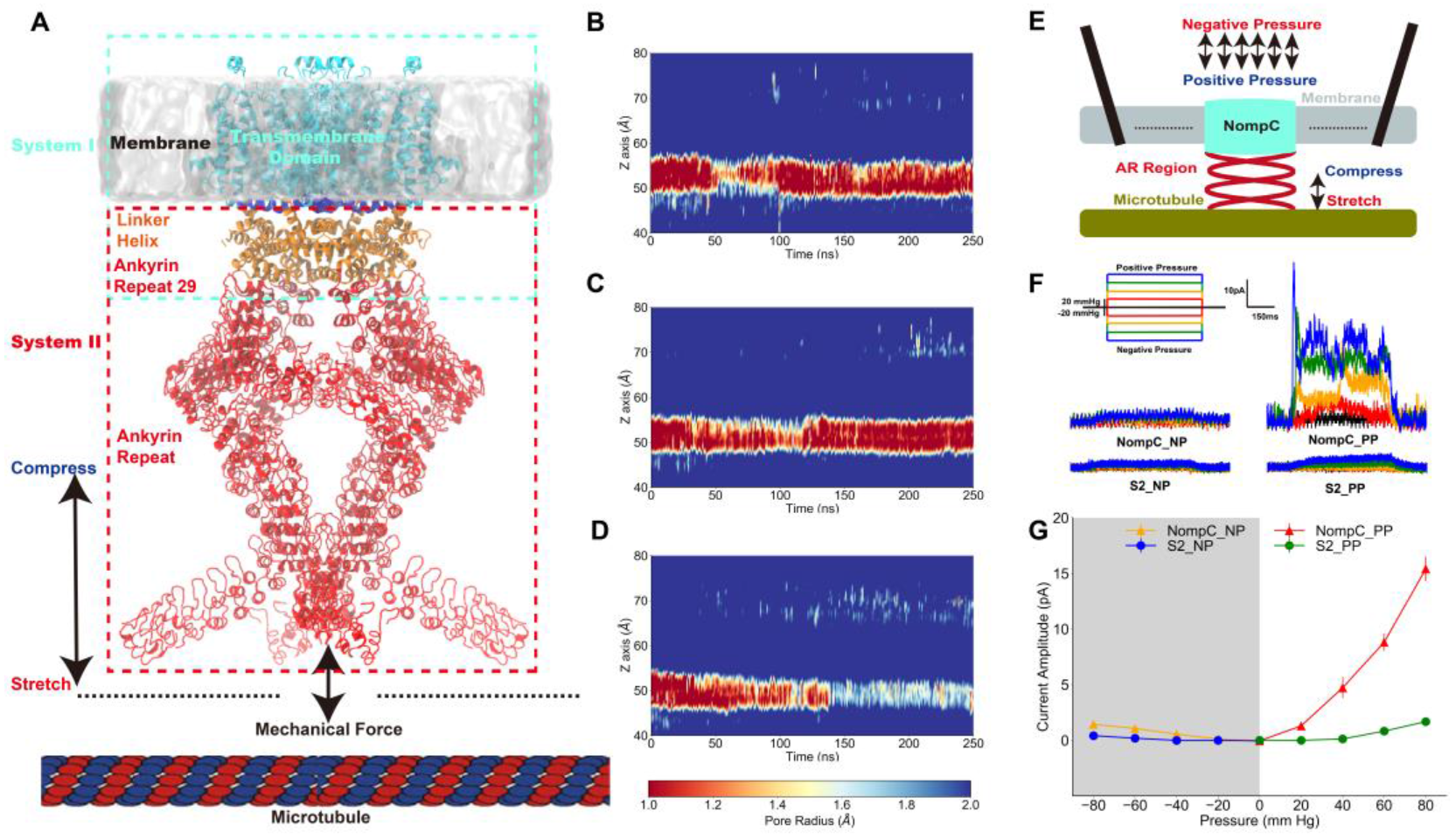
The tethered NompC channel was opened by compression of the intracellular domain. **A.** The simulation systems. The NompC was divided into two subsystems for the molecular dynamics simulations, denoted by the cyan and red rectangular boxes respectively. **B−D.** The TM pore size evolution for the free (**B**), pulling/stretch (**C**), and pushing/compression (**D**) simulations. **E.** A schematic figure of the cell-attached patch-clamp electrophysiological recordings on NompC. **F.** Representative traces of the electrophysiological measurements for the S2 blank cell and NompC-expressed cell, showing that there are significantly larger signals under positive pressure (PP) in the presence of NompC. **G.** The mean and standard deviation of the mechano-gated currents in the S2 blank and NompC-expressed cells under positive and negative pressure (NP) in the cell-attached patch-clamp experiments.

To reveal the atomistic details of how mechanical stimuli can lead to the gating of the tethered NompC channel, we adopted a divide-and-conquer protocol: we performed all-atom molecular dynamics (MD) simulations on the transmembrane and linker helix (TM+LH) domains, and linker helix and ankyrin repeat (LH+AR) domains of the Cryo-EM NompC structure (Fig. 1A). We considered two forms of the most essential forces on the AR helices: pulling and pushing. For the TM+LH system, we applied forces that are normal to the membrane surface on the AR29, which directly connects to the LH region, and monitored how the TM domain responds by calculating the radius of the TM pore. We observed that the channel remains closed (with a very narrow constriction site, radius < 1.0 Å, around the residue I1554) throughout the simulations without any external forces (Fig. 1B), indicating that the closed-state Cryo-EM structure is stable in our “free” MD simulations. When the direction of the pulling force is away from the membrane surface, the TM channel also remains closed in our MD simulations (Fig. 1C). In fact, the narrow region with a radius of less than 1 Å even expanded in the latter part of the trajectory, indicating that the channel was actually more closed than the free NompC in our “pulling” simulations. In contrast, when applying a pushing force (toward the membrane) on the AR29, we observed that the narrowest constriction site of the channel was significantly dilated in the latter part of the “pushing” simulation (Fig. 1D). Moreover, when applying a membrane potential of 300 mV, we observed continuous ion permeation through the dilated pore caused by the pushing force in our MD simulations (Fig. S2-S4). Therefore, our simulation results indicated that the NompC channel may be opened by a pushing force from the intracellular side, but not by a pulling force.

To validate the above findings, we did cell-attached patch-clamp experiments (Fig. 1E), in which positive or negative pressure with a 20 mm Hg increment was applied. Since the AR region was shown to be associated with microtubules (*27*), it was conceivable that a positive pressure will result in a slight compression of the AR region and thus a pushing force on the TM domain, whereas a negative pressure will generate a slight stretch of the AR region and a pulling force on the TM domain in the cell-attached patch-clamp experiments. As shown in Fig. 1F and G, the reference *Drosophila* S2 cells without NompC expressed showed no response to the positive and negative pressure stimuli, while we can detect a clear electrical signal through the NompC-expressed S2 cells under a positive pressure, whereas the signal under negative pressure is nearly negligible. Previous studies have shown that the AR regions are necessary for the mechano-gating of NompC, and the force from lipids cannot open the channel in the absence of the AR region (*14*). Therefore, our experimental results indicated that it is the compression of the AR region, and the resulted pushing force, that opens the channel, which is consistent with our MD simulations.

We then investigated how a pushing force from the AR region can open the TM pore by analyzing the TM+LH simulations. The free, pulling and pushing trajectories were concatenated, and the principal component analysis (PCA) was performed to visualize the collective motion of the NompC pore domain. As shown in Fig. 2A, the second PCA eigenvector can distinguish the conformations of the free, pushing and pulling simulations very well, with the larger values corresponding to the more dilated states. We extracted the two extreme conformations along the second PCA eigenvector and overlaid them to visualize the most significant conformational changes of the TM domain under the three mechanical stimuli (Fig. 2B). We observed an evident clockwise rotation of the TRP domain when a pushing force was applied to the AR29 (looking from the intracellular side, Fig. 2B, Fig. S7 to S8). Such clockwise rotation of the TRP domain was proposed to be associated with the opening of the TRP channels (*28*–*31*). Indeed, the overlaid structure in Fig. 2B showed that the clockwise rotation of the TRP domain induced the S6 helices (which is directly linked to the TRP domain) to rotate clockwise as well, albeit to a lesser extent. The gating constriction site was found to be located at I1554 of S6 helix (*18*), and in our simulations, they were pulled away from the channel axis when the S6 helices rotate clockwise together with the TRP domain, leading to the dilation of the pore (Fig. 2B and C). Thus, consistent with previous structural studies of TRPV1 (Fig. S9) (*28*–*30*), our simulations showed that the clockwise rotation of the TRP domain (as well as the S6 helices) leads to the opening of the NompC pore. And it is the pushing force (compression of the intracellular domain) that leads to such a collective gating motion.

**Figure 2.**
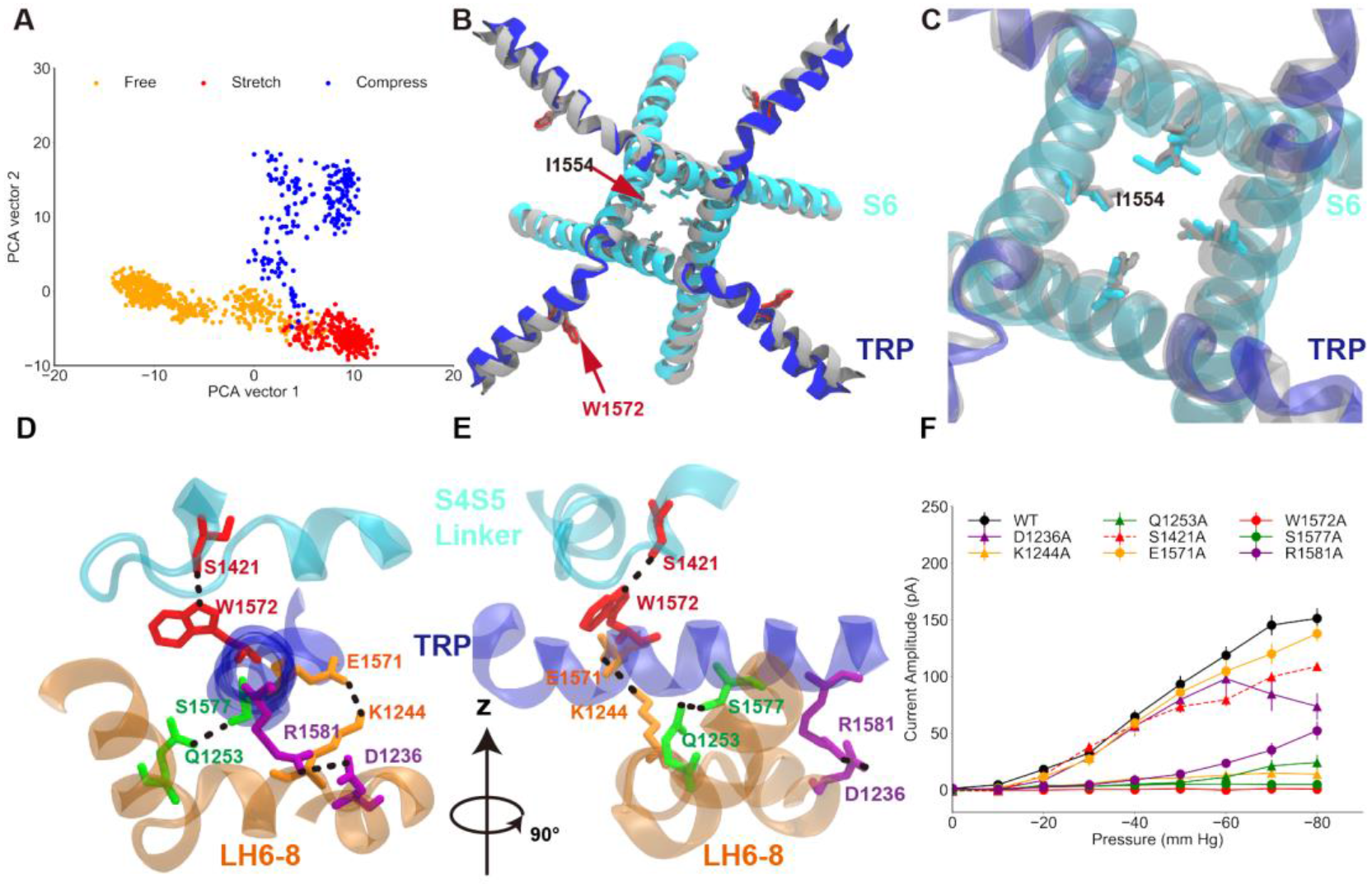
The conformational changes of the TRP and TM domains during gating. **A.** The principal component analysis (PCA) of the MD simulation trajectories. The projections on the second eigenvector can well distinguish the conformations under pulling or pushing. **B.** The overlaid extreme structures along the second eigenvector of the PCA. The most closed conformation (silver) and open conformation (cyan) showed the global changes of the TRP domain during gating: a clockwise rotation. **C.** The orientation and position of the gate residue, I1554, in the most closed (silver) and open (cyan) conformations in the simulations. **D and E.** The residues forming four stable hydrogen bonds between the TRP and LH domains. **F.** The mean and standard deviation of the mechano-gated current of the wild-type NompC, as well as the mutants W1572A, S1421A, Q1253A, S1577A, K1244A, E1571A, D1236A, and R1581A, under negative pressure in the outside-out patch-clamp experiments.

We analyzed the hydrogen bond network around the TRP domain, trying to locate the key residues ensuring the clockwise rotation of the TRP domain in response to the pushing force from AR. We identified four stable hydrogen bonds throughout the MD simulations (Fig. 2D and E and Table S4). These stable hydrogen bonds indicate a conservative interaction network as well as a stable local configuration during the gating process. We then did mutations on the residues forming these hydrogen bonds and performed electrophysiological experiments to check if any of them play crucial roles in the gating of NompC. As shown in Fig. 2F, mutations of most of the eight residues led to loss-of-function to some degree. In particular, the W1572A mutation completely abolished the gating behavior of the channel, in agreement with Jin et al.’s early work (*18*). Interestingly, we found W1572 to be the rotation pivot of the TRP domain in our MD trajectory, which forms a stable hydrogen bond with the backbone of S1421. In addition, the mutations S1577A and R1581A on the TRP domain, and K1244A and Q1253A on the LH domain, all resulted in a significant loss-of-function, indicating the essential roles of these residues in conveying the forces from the AR region to the TRP domain. Thus, five out of seven residues, whose side chains form hydrogen bonds between the TRP and LH domains bonds as identified in our MD simulations, were proven to be crucial for the proper gating behavior of the NompC channel. The other two residues, D1236 and E1571, which are also involved in the hydrogen bonding between the TRP and LH domains, were found to be replaceable by adjacent residues in stabilizing the local conformation (Fig. S10). Taken together, we conclude that W1572 acts as a rotation pivot, while the TRP domain senses a pushing force from the LHs upon AR compression, which was stabilized by at least four hydrogen bonds, resulting in a clockwise rotation of the TRP domain around W1572. This also confirms the notion that a force/conformational change has to be transferred from the AR region to the TRP domain through the LHs when the NompC channel is opening in response to a mechanical stimulus.

To study how a pushing/pulling force is transferred to the LHs from the ARs, we performed multiple MD simulations on the truncated LH and AR domains (Fig. 3A). We applied position restraints on the LHs (orange) and ran simulations with or without external forces applied to the terminal AR1 (Fig. 3A). Several interesting mechanical properties were obtained from these simulations. First, we analyzed the reaction forces of the position restraints on the LHs, which were identical to the forces acting on the LHs by ARs, in terms of magnitude and direction. Our analysis indicated that when pushing the AR1 toward the membrane with a total force of 20 pN, that is 5 pN of force on each chain, the AR29 will apply a total torque of ~13 pN·nm on the LH domain, pointing to the extracellular side, which would help to rotate the TRP domain clockwise and drive the channel to open (Fig. 3B). This is in line with the recent continuum mechanics study by Argudo et al (*21*). Second, we calculated the force constant of the AR spring by 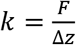, where F is the force we applied on AR1, and ∆*z* is the distortion of the AR region (Fig. 3C). The spring constant of each AR helix was estimated to be 3.3±0.9 pN/nm in the supercoiled helix formed by the four AR chains. For comparison, previous atomic force microscopy measurement obtained a force constant of 1.87±0.31pN/nm (*32*), and early steered molecular dynamics simulations obtained a value of ~4.0 pN/nm (*33*). It should be noted that our calculation was performed for the supercoiled AR helix complex, while the previous study was for a single 24-AR spring. For comparison, we performed SMD on a single 29-AR spring, and estimated the spring constant to be around 2.5±0.4 pN/nm (Fig. S12). The close agreement of the values from the single AR and AR complex indicated that the four AR helices are not tightly coupled. Third, we analyzed how fast the forces can be transferred from the AR1 to the LH. The deviation of the directions of the forces exerting on the LH regions when the AR region was stretched or compressed occurred at the time of about 7~8 ps (Fig. 3D, Fig. S13). Considering the fact that the length of the relaxed AR region is about 15 nm, we estimate that the force was transducted through the AR region at a speed of 1.8 ± 0.2 nm/ps. Interestingly, a recent study showed that the force is propagated via the membrane at a speed of 1.4 ± 0.5 nm/ps (*34*). Therefore, it appears that the speed of force transduction in the tethered NompC channel is faster than that in the membranes.

**Figure 3.**
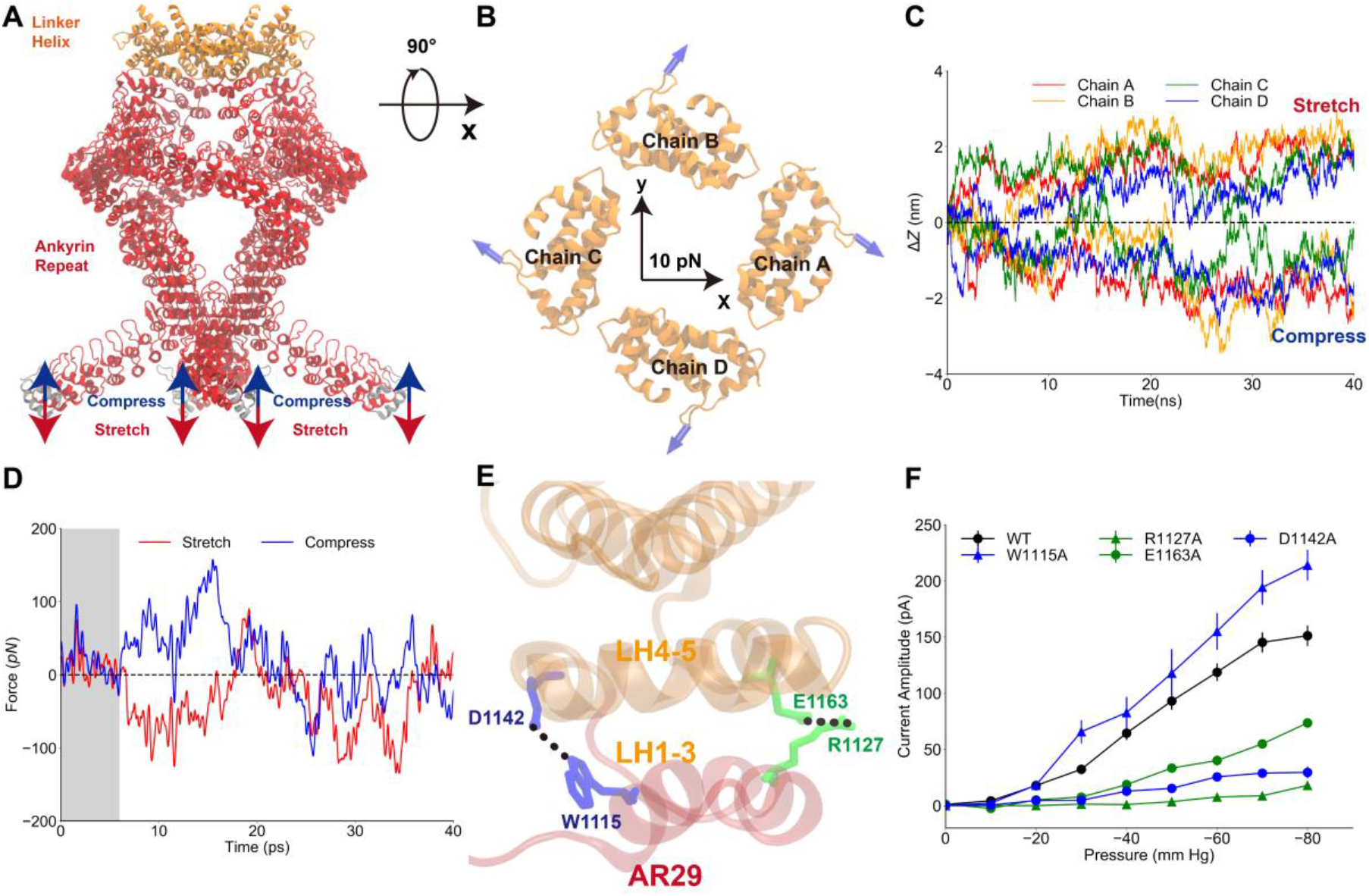
The mechanical properties of the AR region. **A.** The simulation system, in which the LH domain (orange) was restrained and a pushing or pulling force was applied to the first AR (grey). **B.** Projection of the reaction forces of the restraints on the LH domain (same as the forces exerted on the TRP domain by the LH domain) on the plane parallel to the membrane surface, showing that a torque is generated that will drive the LH and TRP domain to rotate clockwise (looking from the intracellular side). The calculation was symmetrized from the original data as shown in Fig. S11. **C.** The AR region was compressed/stretched by a pushing/pulling force of 5 pN and reached its equilibrium length within 40-ns simulations. **D.** The evolution of the reaction forces of the restraints on the LH domain during the simulation, when pushing (blue) or pulling forces (red) were applied to AR1. A clear derivation occurred at around 7~8 ps during the simulation time, indicating that the forces applied on AR1 have reached LH at the time. **E**. The residues forming two stable hydrogen bonds between the LH domain and AR29. **F.** The mean and standard deviation of the mechano-gated current of the wild-type NompC and the mutants W1115A, D1142A, R1127A, and E1163A, under negative pressure in the outside-out patch-clamp experiments.

In addition, we found two stable hydrogen bonds between the ARs and LHs in our MD trajectories, between W1115 and D1142, and R1127 and E1163, respectively (Fig. 3E). Mutations of D1142A, R1127A, and E1163A, which break the hydrogen bonds, lead to a significant loss-of-function. However, W1115A does not alter the mechanosensing behavior very much (Fig. 3F), as its hydrogen bonding and stabilizing role can be replaced by the adjacent Y1109, which can also form a stable hydrogen bond with D1142 in the presence of the mutation W1115A (Fig. S10). These results indicate that the interface between the AR and LH regions is also crucial for the force transduction, further confirming the tethered spring model for NompC.

In summary, our combination study of MD simulations and electrophysiological experiments revealed a clear “push-to-open” gating model of the NompC channel. As illustrated in Fig. 4, when the AR spring is compressed, which can be caused by the compression or shrinking of cells, for instance, the AR spring of NompC will generate not only a pushing force but also a torque on the TRP domain with a direction pointing to the extracellular side and perpendicular to the membrane surface. The torque is generated by the specific supercoiled structure of the AR region, as revealed by the recent continuum mechanics study of Argudo et al. as well (*21*). This torque will drive the TRP domain to rotate clockwise. Intriguingly, our simulations indicate that even the pushing force itself is probably enough to generate a clockwise motion of the TRP domain, which in turn pulled the S6 helices to open the gate of the NompC. A few critical residues between the TRP domain and LH, including R1581, W1572, Q1253, S1577 and K1244, as well as the presence of the S4-S5 linker above the TRP domain, ensures that the TRP domain will rotate clockwise around the pivot W1572 when a pushing force is applied on the LH region. This is supported by a recent study showing that a mutant of TRPV1, which has only two ARs, can be mechanically opened by a pushing force as well (*35*). Therefore, we hypothesize that the TRP channels similar to NompC, with a certain number of ARs, can be tethered to microtubules and utilize the push(-AR)-to-open mechanism to feel and respond to cell shrinking or compression, which can be a complementary sensing mechanism to the well-studied stretch(-membrane)-to-open mechanism that responds to cell expansion.

**Figure 4.**
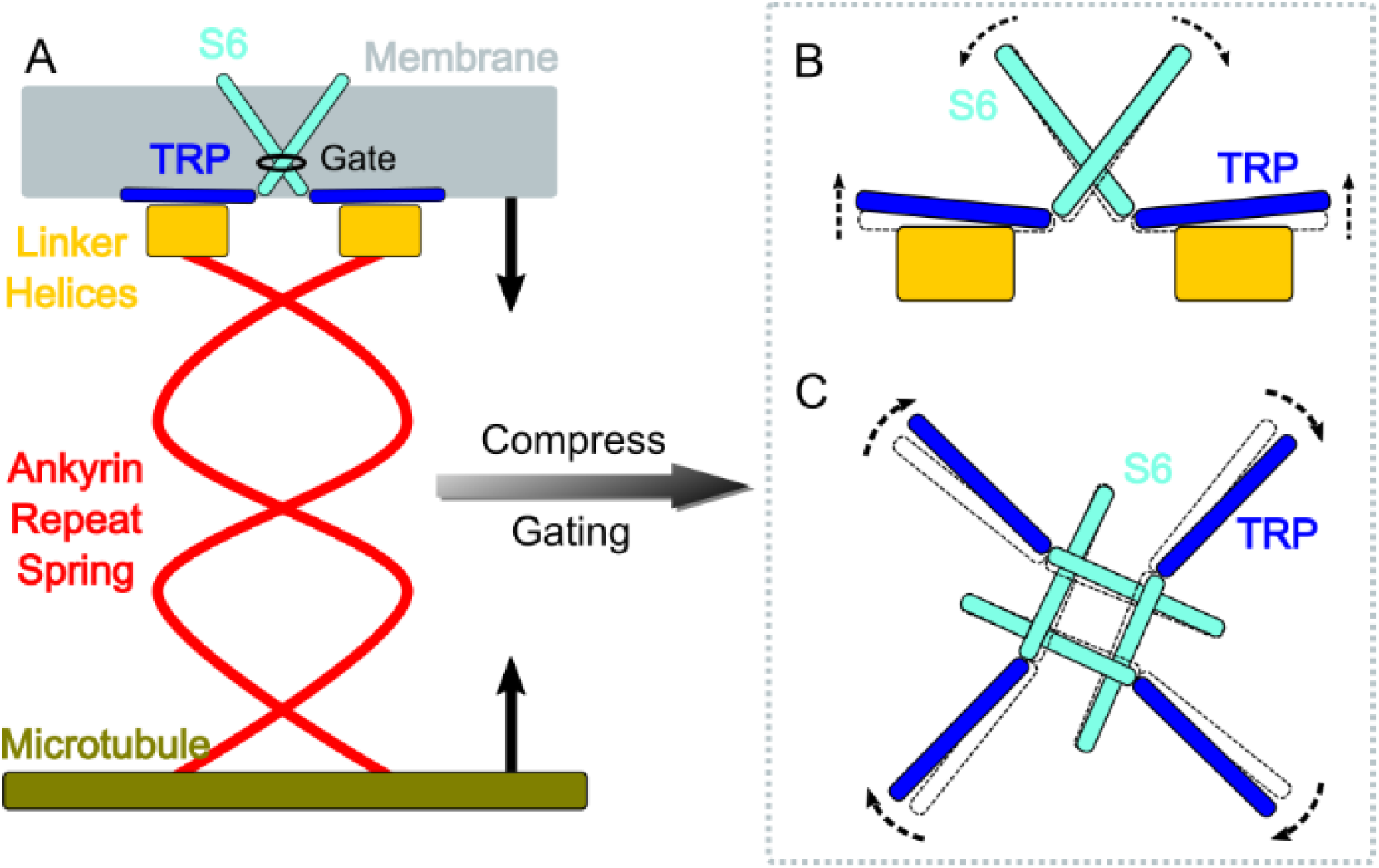
A sketched gating model of NompC. **A.** The compression of the AR region will generate a pushing force and a torque on the LH domain, pointing to the extracellular side. **B.** The LH domain further pushes the TRP domain, leading to a tilt (side view), and **C.** a clockwise rotation of the TRP domain (looking from the intracellular side). The motion of the TRP domain pulls the S6 helices to slightly tilt and rotate, which dilates the constriction site of the pore.

## Acknowledgment

The research was supported by the National Key Research & Development Program of the Ministry of Science and Technology of China (2016YFA0500401 to C.S., and 2017YFA0103900 & 2016YFA0502800 to Z.Y.), and the National Natural Science Foundation of China (21873006, 31571083 and 31970931). Z.Y. was supported by the Program for Professor of Special Appointment (Eastern Scholar of Shanghai, TP2014008), the Shanghai Municipal Science and Technology Major Project (No.2018SHZDZX01) and ZJLab, and the Shanghai Rising-Star Program (14QA1400800). Part of the molecular dynamics simulation was performed on the Computing Platform of the Center for Life Sciences at Peking University, and part of the molecular dynamics simulation was performed on the Tianhe II supercomputer in the National Supercomputing Center in Tianjin.

