## Supplementary Information for "Push-to-open: The Gating Mechanism of the Tethered Mechanosensitive Ion Channel NompC"

[1]State Key Laboratory of Medical Neurobiology and MOE Frontiers Center for Brain Science, Human Phenome Institute, Ministry of Education Key Laboratory of Contemporary Anthropology, Collaborative Innovation Center of Genetics and Development, Institute of Brain Science, Department of Physiology and Biophysics, School of Life Sciences, Fudan University, 2005 Songhu Road, Yangpu District, Shanghai 200438, China

[2]Center for Quantitative Biology, Academy for Advanced Interdisciplinary Studies, Peking University, Beijing 100871, China

[3]Peking-Tsinghua Center for Life Sciences, Academy for Advanced Interdisciplinary Studies, Peking University, Beijing 100871, China

[#]These authors contributed equally to this work.

### S1. Methods and Material

#### S1.1 The simulation systems

We adopted a “divide-and-conquer” strategy for the molecular dynamics (MD) simulations, and simulated two systems separately. System I includes the transmembrane (TM) region, the linker helix (LH) region and the ankyrin repeat (AR) 29 of NompC. The PPM server was used to reorient the NompC structure to ensure that the TM domain of NompC was well located in a lipid bilayer (*1*). The protein was embedded in a 1-palmitoyl-2-oleoyl-sn-glycero-3-phosphocholine (POPC) bilayer and then solvated in a water box of 150×150×150 Å$^3$. CHARMM-GUI was used to generate the configuration and topology of the simulation system, as well as the parameter files with the CHARMM36m force field (*2-4*). There were 492 POPC molecules, 72,000 water molecules, and sodium and chloride ions corresponding to a concentration of 150 mM in the setup, resulting in a system of 314,352 atoms in total.

System II includes the LH domain and the AR domain of NompC. The protein was solvated in a water box of 200×200×200 Å$^3$. CHARMM-GUI was used to generate the configuration, topology and parameter files with CHARMM36m force fields. In addition to the protein, 354,567 water molecules were added and sodium chloride ions were added to maintain an ion concentration of 150 mM. The simulation system II contains 1,134,213 atoms in total.

#### S1.2 Molecular dynamics simulations

All the MD simulations were performed with GROMACS 5.1.2 (*5*). The REDUCE program in AMBER was used to add hydrogens to the original PDB files and determine the protonation state of the histidine residues (*6, 7*). For the system I, energy minimizations were achieved using the steepest descent algorithm, followed by a two-stage equilibration, a 0.4-ns NVT (constant particle number, volume, and temperature) equilibration simulation with harmonic restraint applied to the protein molecules (a force constant of 4000 kJ mol$^{-1}$nm$^{-2}$ on the backbone and 2000 kJ mol$^{-1}$nm$^{-2}$ on the side chains), and a 20-ns NPT equilibration simulation with gradually decreased restraint (from 2000 kJ mol$^{-1}$nm$^{-2}$ to 100 kJ mol$^{-1}$nm$^{-2}$ on the backbone and from 1000 kJ mol$^{-1}$ nm$^{-2}$ to 50 kJ mol$^{-1}$nm$^{-2}$ on the side chains). During the equilibration processes, harmonic restraints were applied to heavy atoms of the protein and planar restraints were used to keep the positions of lipid head groups along the normal direction of the membranes. The simulation temperature of the system was set to 300 K. Once all the equilibration steps were completed, the restraints were removed and the production simulations were performed in the NPT ensemble. The time step was 2 fs. The cubic periodic boundary condition was used during the simulations and the van der Waals interaction cut-off was set to 12 Å. The long-range electrostatic interactions were calculated with the Particle Mesh Ewald (PME) method (*8*).

For the system II, the steepest descent algorithm was used to achieve initial energy minimizations and then followed by a two-stage equilibration, a 0.2-ns NVT (constant particle number, volume and temperature) equilibration simulation with harmonic restraint forces applied to the protein (force constants of 400 kJ mol$^{-1}$nm$^{-2}$ on the backbone and 40 kJ mol$^{-1}$nm$^{-2}$ on the side chains), and a 10-ns NPT equilibration simulation with restraints on the protein backbone (force constant of 400 kJ mol$^{-1}$nm$^{-2}$ ) and side chains (force constant of 40 kJ mol$^{-1}$nm$^{-2}$). The temperature was set to 300 K. In the production simulations of the system II, 1000-kJ mol $^{-1}$nm$^{-2}$ harmonic restraints were applied to the heavy atoms of the LH domain while the restraints on the AR region were removed. Multiple 40-ns trajectories were generated in the NPT ensemble with a time step of 2 fs. The trajectories were saved every 10 ps. The long-range electrostatic interactions were calculated with the PME method (*8*).

#### S1.3 Steered molecular dynamics simulations

For the system I, after equilibration, steered molecular dynamics (SMD) was utilized to pull AR29 to simulate the mechanical stimuli from the AR region (spring) (*9*, *10*). The TM regions of the four chains of NompC were treated as the reference groups, while the AR29 of the four chains were treated as the pulling groups. In this work, in addition to free simulations (no pulling forces on the AR region), we considered the two most essential mechanical stimuli: the pulling and pushing forces on the AR29 along the direction normal to the membrane surface (the z-axis in our simulations), where pulling meaning the force is pointing to the intracellular side (stretch of the AR spring) and pushing meaning the force is pointing to the extracellular side (compression of the AR spring) along the z-axis. We tested a series of harmonic force constants (from 50 kJ $mol^{-1}nm^{-2}$ to 200 kJ $mol^{-1}nm^{-2}$) to pull AR29. The pulling speed of AR 29 was as slow as 0.1 Å/ns in order to leave sufficient time for the TM region of NompC to relax. During the MD/SMD simulations, the distances between the TM region of NompC and AR29, and the driving forces that act on the four chains of AR29 were recorded. The frames from MD/SMD trajectories were saved every 1 ns. All the MD/SMD trajectories of the system I are listed in table S1.

**Table S1**. MD/SMD trajectories of system I

| Trajectory | Pulling Group | Force Constant ($kJ\cdot mol^{-1}nm^{-2}$) | Pulling or Pushing | Simulation Time (ns) |
|---|---|---|---|---|
| Free I | / | 0 | / | 500 |
| Compress I 1 | AR29 | 50 | pushing | 250 |
| Compress I 2 | AR29 | 100 | pushing | 250 |
| Compress I 3 | AR29 | 200 | pushing | 250 |
| Stretch I 1 | AR29 | 50 | pulling | 250 |
| Stretch I 2 | AR29 | 100 | pulling | 250 |
| Stretch I 3 | AR29 | 200 | pulling | 250 |

For the system II, starting from the equilibrated structure, the LH domain was position-restrained and SMD simulations were performed to pull AR1, simulating the compression and stretch of the AR region (*9, 10*). The LH domains of the four chains of NompC were treated as the reference group, while the AR1 of the four chains were treated as the pulling groups. Constant pulling forces of 5 pN were applied on AR1 of each chain along the z-axis. Again, we considered the two most essential mechanical stimuli here: the pulling and pushing forces on the AR1, where pulling meaning the force is pointing to the intracellular side (stretch of the AR spring) and pushing meaning the force is pointing to the extracellular side (compression of the AR spring) along the z-axis. During the SMD simulations, the distances between the LH domain and the AR1 of each chain were recorded. To take into account the position restraints applied to the AR1 of the four chains by microtubules, an additional flat bottom potential of 100 kJ $mol^{-1}nm^{-2}$ with a radius of 3 nm was added on the four chains of AR1 on the x-y plane, to restrain each AR1 to move within a cylinder parallel to the z-axis. To estimate the mechanical property of a single chain of AR, the same protocol was applied to the single chain A of system II.

To estimate the speed of the forces that convey along the AR spring, five 40-ps trajectories (Free 2-6, Compress 2-6 and Stretch 2-6 in table S2) were generated for each condition (free, compress/push and stretch/pull) with a constant force of 5 pN on AR1 of each chain. The frames from trajectories were saved every 10 fs, a frequency high enough for the force transfer analysis. All the MD/SMD trajectories of system II are listed in table S2.

**Table S2**. MD/SMD trajectories of system II.

| Trajectory | Pulling Group | Force (pN) | Pulling or Pushing | Simulation Time (ns) |
| --- | --- | --- | --- | --- |
| Free II 1 | / | / | / | 40 |
| Compress II 1 | AR1 | 5 | pushing | 40 |
| Stretch II 1 | AR1 | 5 | pulling | 40 |
| Free II 2-6 | / | / | / | 0.04 × 5 |
| Compress II 2-6 | AR1 | 5 | pushing | 0.04 × 5 |
| Stretch II 2-6 | AR1 | 5 | pulling | 0.04 × 5 |
| Free II 7 Chain A | AR1 Chain A | / | / | 100 |
| Compress II 7 Chain A | AR1 Chain A | 5 | pushing | 100× 3 |
| Stretch II 7 Chain A | AR1 Chain A | 5 | pulling | 100× 3 |

#### S1.4 The ion permeation simulations

After 200-ns simulations for the system I, the TM pore of NompC was partially opened (Fig. 1). As can be seen in Fig. S1A, removing the pushing force, the partially opened structure became closed again. This partially open state can be maintained by an umbrella pushing potential with a force constant of 100 kJ $mol^{-1}nm^{-2}$ and an initial force of 50 pN on the AR29 of each chain. The open state of NompC ion channel was further simulated for more than 500 ns, and a transient structure with the pore radius of the lower constriction more than 2.0 Å was obtained at 545 ns (Fig. S1B). This structure was wide enough for partially hydrated cations to permeate and therefore was used to simulate the ion permeation processes. The gate region, which include S5, S6, the selectivity filter and TRP region, was position restrained with the harmonic potential with a force constant of 1000 kJ $mol^{-1}nm^{-2}$, while a transmembrane potential of 300 mV was applied (by setting a uniform electric field along the z-direction) to drive the ions to pass through the channel. Three independent 200-ns MD trajectories were generated with 150-mM KCl or NaCl in the systems, respectively. The pore was constantly dilated in these simulations (Fig. S2). The trajectories were saved every 50 ps and the number of ions that permeated through the channel was counted (Fig. S3 and Fig. S4). From the ion permeation count, we calculated the current by $I = \Delta q/\Delta t$, which was then used to calculate the conductance by $C = I/U$, where $U$ was the transmembrane potential 300 mV. The estimated conductance of the channel was about 7~15 pS for the monovalent cations. These values are significantly smaller than those obtained in electrophysiological experiments, so we believe that the dilated NompC structure from our pushing simulations is only partially open. Unfortunately, it is still highly challenging to simulate a fully-opened structure purely by MD simulations, and it is beyond the scope of the present work.

**Table S3**. The ion permeation simulations of the partially open NompC.

| Trajectory | Transmembrane Potential (mV) | Simulation Time (ns) |
|---|---|---|
| Partially Open NompC | 300 | 200 × 3 |

##### S1.5 The principal component analysis (PCA)

For the MD simulations of system I without any external forces on the AR29, the distance between the center of mass (COM) of the TM domain and AR29 of NompC (which we term "TM-AR29 distance" here) fluctuated around 20 Å for each chain in the MD trajectories (Fig. S5A), indicating that the overall structure of NompC remained stable under the "free" condition. For the MD trajectory representing the stretch of ARs (pulling AR29 toward the intracellular side), the TM-AR29 distance slowly increased during the first stage of the simulation **(**Fig. S5B**)**. However, after 250 ns (the light gray area in Fig. S5B), the TM-AR29 distance of Chain D exhibited a sharp increase, which represented the secondary structure of chain D being disrupted at this time period. Therefore, the frames of trajectory after 250 ns were disregarded for further PCA analysis. For the MD simulations that represent the compression of the AR domain (pulling the AR29 toward the extracellular side, or "pushing"), the TM-AR29 distance was gradually decreased (Fig. S5C). After 200 ns (black dashed line in Fig. S5C), the TM-AR29 distance of Chain D became significantly shorter than the other three chains, indicating that the symmetry of the NompC structure was broken due to the excessive pushing force disrupting chain D during this time. Therefore the 200-ns structure was treated as the final structure of the pushing SMD trajectory. Correspondingly, the 200-ns structure of the free MD trajectory and 200-ns structure of the pulling SMD trajectory were selected as the final structures for comparison, as shown in Fig. S6.

The frames after 200 ns of the pulling trajectory were disregarded for the PCA analysis. The 950 protein structures (500 frames from the free MD trajectory, 250 frames from the stretch/pulling SMD trajectory and 200 frames from the compress/pushing SMD trajectory) were concatenated for the PCA analysis, as shown in Fig. 2A. The second eigenvector can well distinguish the structures from the free, pushing or pulling simulations. Moreover, as shown in Fig. S7A, the pore size of the NompC structure projected along the second eigenvector showed a clear trend of opening at the lower gate for the "pushing" simulation. Therefore, we think the second eigenvector of PCA may represent the conformational change of gating.

#### S1.6 Analysis of the motion of the TRP domain

The TRP domain of NompC showed a slight tilt (side view) and a clear clockwise rotation (bottom view) upon the opening of the NompC. The tilt angle was defined as the angle between the z-axis and the axis of the TRP alpha helix. As shown in Fig. S7B**,** the tilt angle of the TRP domain was increased from about 0° (frame 0 to 800) to a value of about 2° (frame 800 to 950). More importantly, the TRP domain of NompC showed an evident clockwise rotation (looking from the intracellular side). As shown in Fig. S7C, the rotation angle of TRP domain was increased from about 0° (frame 0 to 800) to an average value of about 6° (frame 800 to 950). The tilt and clockwise rotation of the TRP domains of the four chains were highly correlated with the opening of the lower gate (Fig. S7A), indicating that these motions of the TRP domain were necessary for the gating of NompC.

We also analyzed the evolution of the average tilt angle and rotation angle of the four chains of the TRP domain in the pulling/pushing SMD trajectories with respect to the equilibrated structure (the 200 ns structure obtained from the free MD trajectory), as shown in Fig. S8. Despite the motion of the four chains being asymmetric, the average tilt and rotation angle of them showed a similar tendency as the PCA results indicated. As shown in Fig. S8A and Fig. S8B, the TRP domain of NompC from the final structure of the pulling SMD trajectory showed slight tilt toward the intracellular side and clear counter-clockwise rotation in the bottom view. On the other hand, as shown in Fig. S8C and Fig. S8D, the TRP domain of the final structure of the pushing SMD trajectory showed slight tilt toward the extracellular side and evident clockwise rotation in the bottom view. The evolution of the average tilt angle of the TRP domain of the four chains is shown in Fig. S8E. The evolution of the average rotation angle of the TRP domain of the four chains is shown in Fig. S8F.

#### S1.7 The analysis of the open- and closed-state structures of TRPV1

Both the open-state and closed-state structures of the TRPV1 channel have been obtained. Since NompC and TRPV1 share high structural similarities, we analyzed how the TRP domain in TRPV1 changes upon opening. As shown in Fig. S9, we overlaid the closed (PDB accession number: 5irz) and open structures (PDB accession number: 5irx) of TRPV1 that solved in 2016 by aligning their S1-S5 TM domains (*11*). From the side view, we can also observe a tilt of the TRP domain. The TRP domain of the open TRPV1 showed a slight tilt angle of about 2° (Fig. S9A). When viewed from the intracellular side (Fig. S9B), the TRP domain of the open TRPV1 structure showed evident clockwise rotation (about 9°) in the x-y plane with respect to the closed structure.

Therefore, the analysis results of the open and closed TRPV1 structures were very similar to our results of NompC from the MD/SMD trajectories (Fig. 2, Fig. S7 and Fig. S8).

#### S1.8 Identification of the critical residues from the MD trajectories

To investigate how the mechanical force caused by the stretch and compression of the AR domain is passed onto the TM domain, we analyzed the stable hydrogen bonds and the salt-bridges between the AR29 and the linker helices (LHs), as well as between the LHs and the TRP domain in the MD/SMD simulation trajectories shown in Fig. 1B to Fig. 1D. The relevant residue pairs that form hydrogen bonds are listed in table S4.

**Table S4:** The stable hydrogen bonds and their occupancies in the MD/SMD trajectories.

| Residue 1 | Residue 2 | Free | Stretch | Compress |
|---|---|---|---|---|
| D1236 (LH) | R1581 (TRP) | 98% | 97% | 98% |
| S1421 (4S5) | W1572 (TRP) | 60% | 81% | 81% |
| Q1253 (LH) | S1577 (TRP) | 56% | 44% | 73% |
| K1244 (LH) | E1571 (TRP) | 94% | 94% | 97% |
| W1115 (AR29) | D1142(LH) | 88% | 91% | 91% |
| R1127 (AR29) | E1163(LH) | 91% | 90% | 90% |

The hydrogen bonds listed in the above table were relatively stable, even in the presence of the external force stimuli. Therefore, we believe these residues may be important for the stability of the local conformation and may play a role in the force conveys through the LH domain and did further mutation experiments and MD simulations to verify. Please also refer to Fig. S10.

##### S1.9 Analyzing the role of the AR region in the force convey

For the tetramer, we calculated the net forces on the LH domain projected in the x-y plane, generated by the compression or stretch of the AR domain (Fig. S11). The average net force in the x-y plane was calculated from 20 ns to 40 ns in the trajectories. Looking from the intracellular side, the compression of the AR domain tends to generate a torque pointing to the extracellular side which can drive the LH domain to rotate clockwise (blue arrows in Fig. S11). On the other hand, the stretch of the ARs domain tends to generate a torque pointing to the intracellular side, which will drive the LH domain to rotate counter-clockwise (red arrows in Fig. S11).

For the single AR chain, we overlaid the final structures of the free (silver), pushing (blue) and pulling (red) simulations after 100-ns SMD/MD simulations (Fig. S12A). As shown in Fig. S12B, the distance between the center of the LH and AR1 (which can be viewed as the length of the AR region) was extended and became stable in 60~100 ns for about 2.0±0.1 nm along the z-axis under stretch or compress. As the applied force was 5 pN, by using the formula $k = F/\Delta z$, we estimated the spring constant of a single AR chain to be about 2.5±0.4 pN/nm (pushing/pulling). The estimated force constant of a single AR chain was slightly larger than the previous AFM experiment results of 1.87±0.31 pN/nm by Lee et al. (*12*) and smaller than the SMD simulation estimation of 4 pN/nm by Sotomayor et al. (*13*). Notably, the spring constant of a single AR chain is smaller than that of the supercoiled helix composed of four AR chains (3.3±0.9 pN/nm), indicating that there is a certain degree of coupling among the four chains when they form a complex.

To estimate the force transfer speed through the AR region, we analyzed how long it takes for the force applied to the AR1 to impact the LH domain (Fig. S13A). We generated five short trajectories (40 ps each) for the free/pushing/pulling simulations of system II with high output frequency (10 fs per frame), as shown in table S2. In the first stage of each trajectory, we find that the force on the LH region cannot be distinguished by the simulation conditions, and then at some point, the force values on the LH domain start to deviate among the free/pushing/pulling simulations (Fig. S12, where grey areas end). This represents the forces applied to AR1 starting to impact the LH domain. Therefore, we estimated that the forces applied on AR1 need about 6~12 ps to arrive at the LH domain, and the speed of force transfer was estimated to be 1.8±0.2nm/ps along the AR region.

#### S1.10 Electrophysiological experiments

Drosophila S2 cells were cultured in Schneider's Insect medium supplied with 10% FBS at 27$^{o}$C. TransIT$^{R}$-Insect Transfection Reagent (Mirus) was used to transfect cells according to the product protocol. pUAST-NompC-EGFP (wildtype or mutants) constructs were co-transfected with pGal4. Recordings were carried out 36-48 hours after transfection.

Electrophysiological recordings were carried out under Olympus CKX41 microscopy equipped with a 40× water immersion lens. Transfected cells were identified by green fluorescence. The sample rate was 10 kHz and filtered at 1 kHz (low-pass). Patch electrodes with 12-20 MΩ resistance were used. The bath solution contains: 140 mM NaMES (sodium methanesulfonate), 10 mM HEPES. For cell-attach mode recording, the pipette solution is the same as the bath solution. For inside-out and outside-out mode recording, the pipette solution contains: 140 mM potassium D-Gluconate, 10 mM HEPES. All solutions were adjusted to 285 mOsm and pH 7.2.

After forming specific recording mode (cell-attach mode, inside-out mode or outside-out mode). Negative pressure or positive pressure was applied to the excised membrane via a high-speed pressure clamp (HSPC, ALA-scientific). Signals generated from pClamp software were sent to HSPC to control the timing and intensity of the pressure.

To record the dose-response curve of the mechanosensitive current, pressure steps of 500 ms with 10 mm Hg (for inside-out and outside-out recording) or 20 mm Hg (for cell-attach recording) were applied to the membrane patch through the recording pipette. The outside-inside-out and outside-out patch-clamp traces under different pressure were shown in Fig. S14A while the mean current under different pressure were shown in Fig. S14B.

#### S1.11 The mutation experiments

All point mutations on NompC plasmid were introduced by site-directed mutagenesis and confirmed via sequencing of the full-length construct. Further experiments were performed the same as outside-out and inside-out patch clamp in wild type NompC.

All point mutations on NompC plasmid were introduced by site-directed mutagenesis using CloneExpress$^{R}$ II One-step Cloning kit (Vazyme) and confirmed via sequencing of the full-length construct. Further experiments were performed the same as outside-out and inside-out patch clamp in wildtype NompC described in S1.10.

#### S3 Supplementary Figures

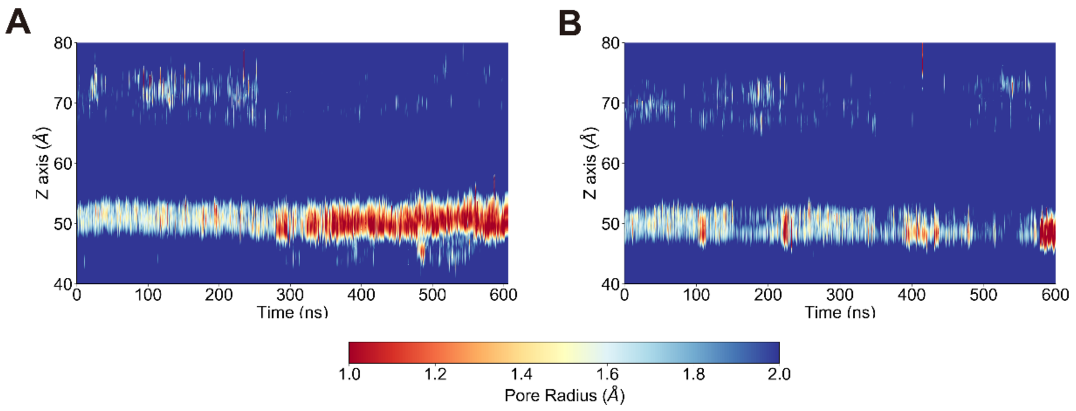


**Figure S1.** The TM pore size evolution for **(A)** the simulation after removing the pushing force, and **(B)** the simulation with a continuous pushing force and a transmembrane potential of 150 mV.

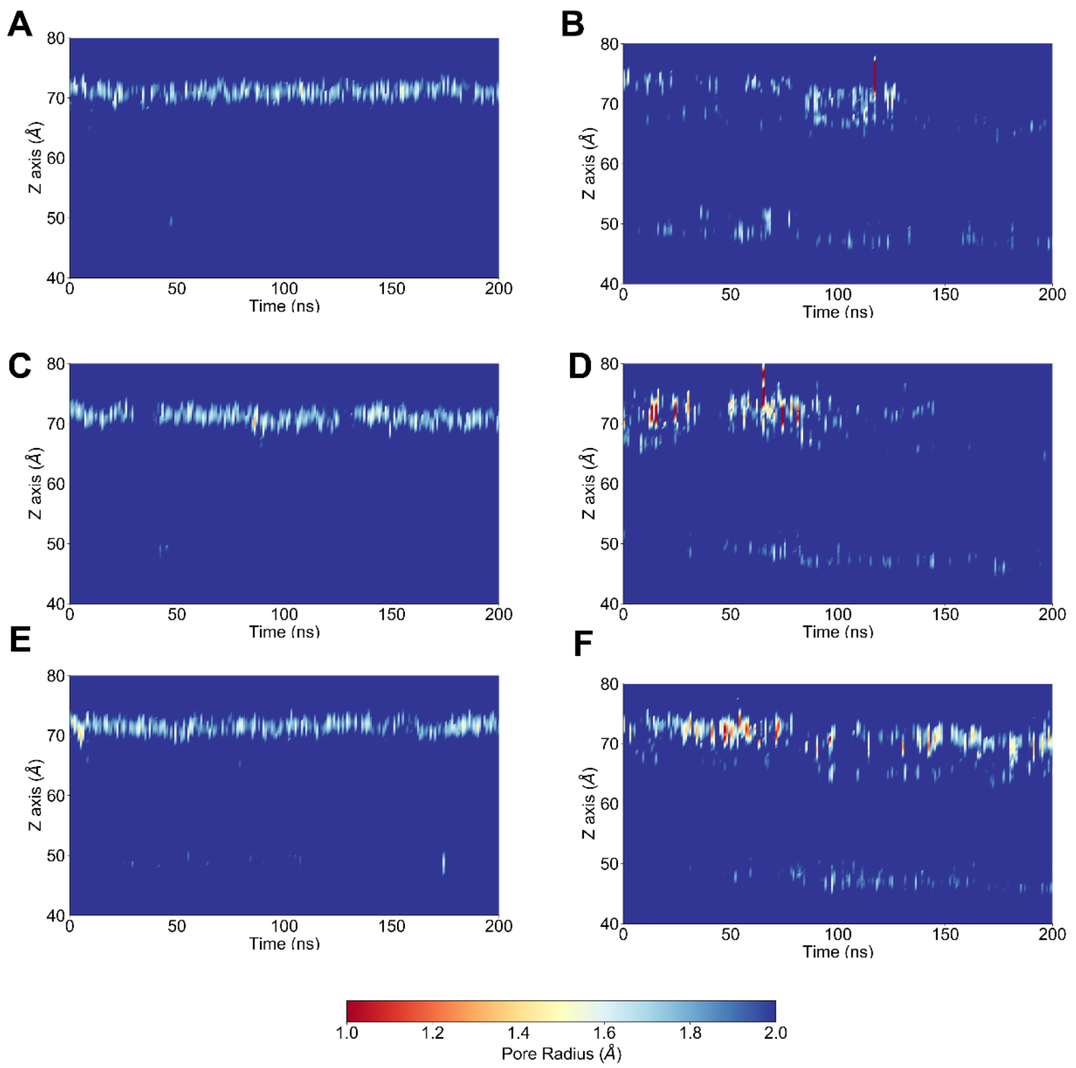


**Figure S2.** The TM pore size evolution in the permeation simulations with NaCl (**A, C, E**) and KCl (**B, D, F**) in the systems.

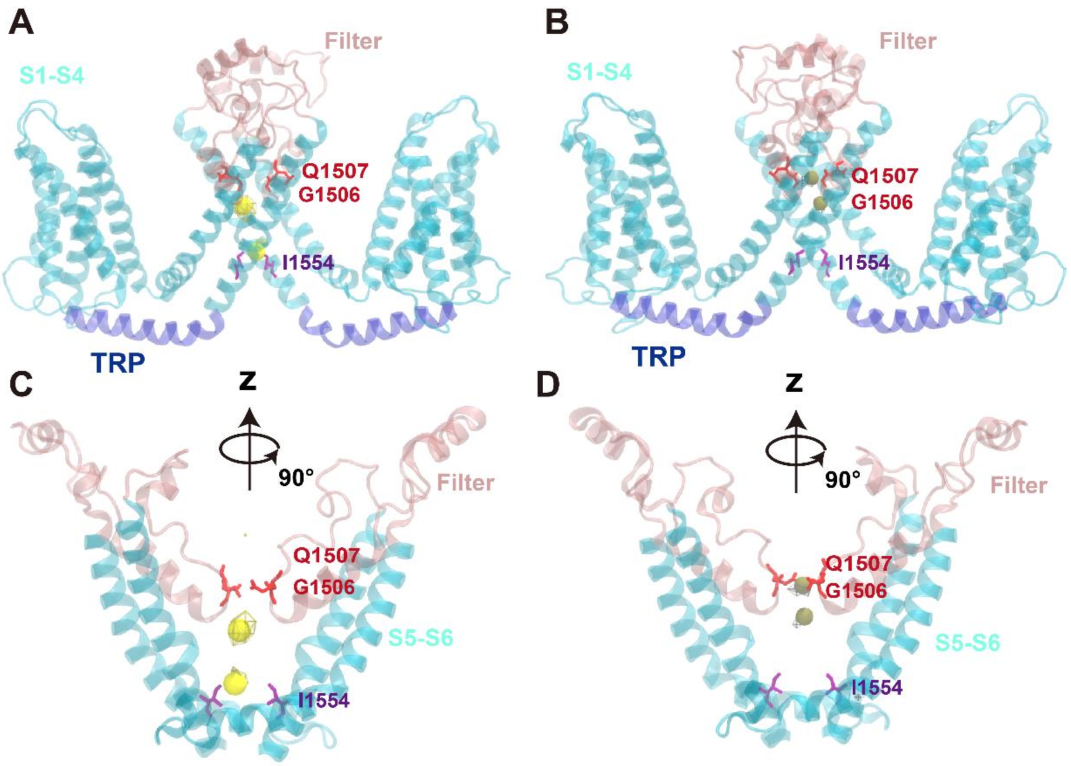


**Figure S3.** The ion density maps from the ion permeation trajectories. The ion density maps were obtained from the simulation trajectories with a -300-mV transmembrane potential for $Na^+$ ions (**A, C**) and a 300-mV transmembrane potential for $K^+$ ions (**B, D**). The TRP domain is colored in blue, transmembrane helices (S1-S6) in cyan and the upper filter in pink. The residues forming the upper gate (G1506 and Q1507) are shown as red sticks and the lower gate (I1554) as purple sticks.

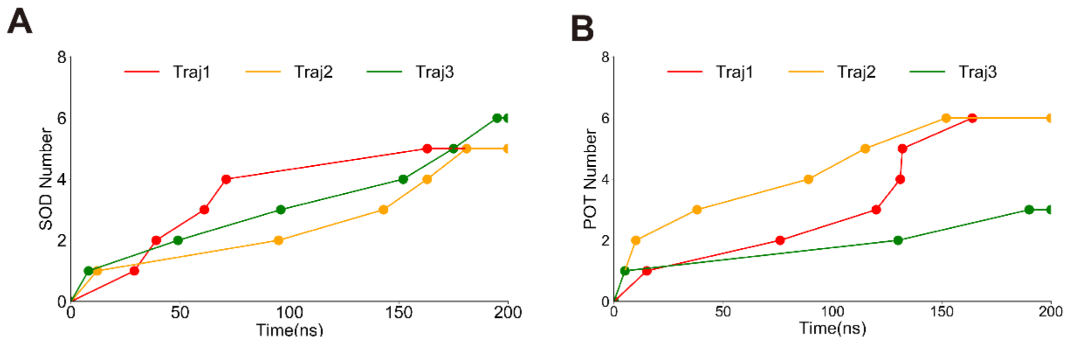


**Figure S4.** Ion permeation count**.** The sodium ion **(A)** and potassium ion (**B)** permeation count through the partially opened structure of NompC in our MD simulations**.**

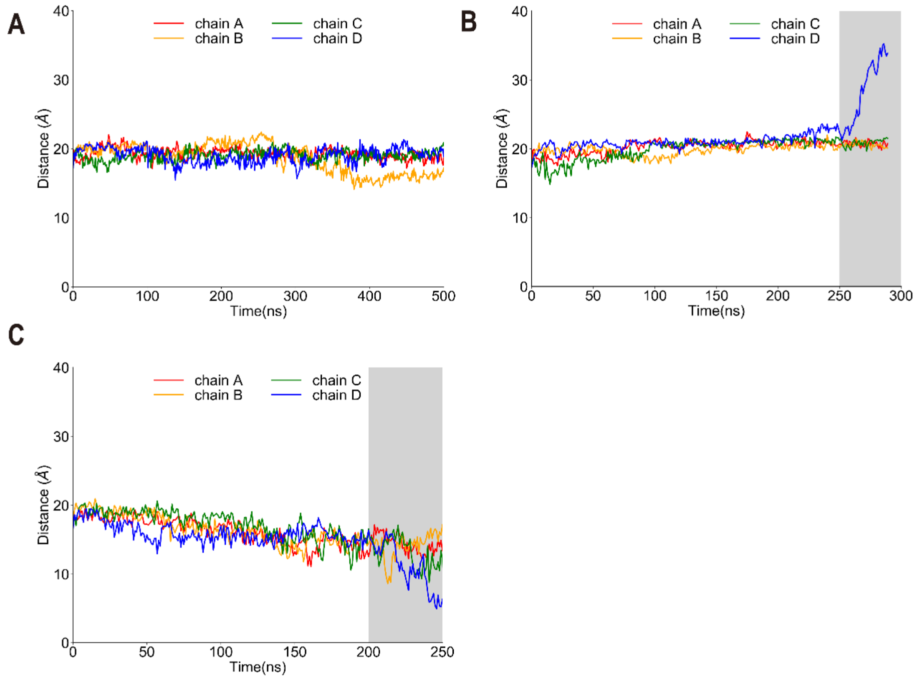


**Figure S5.** Distances between the centers of AR29 and the TM domain of NompC in the MD/SMD simulations of the system I: **A** (free), **B** (pulling), **C** (pushing). The overall structure of NompC remains stable in the white area but gets distorted in the grey area.

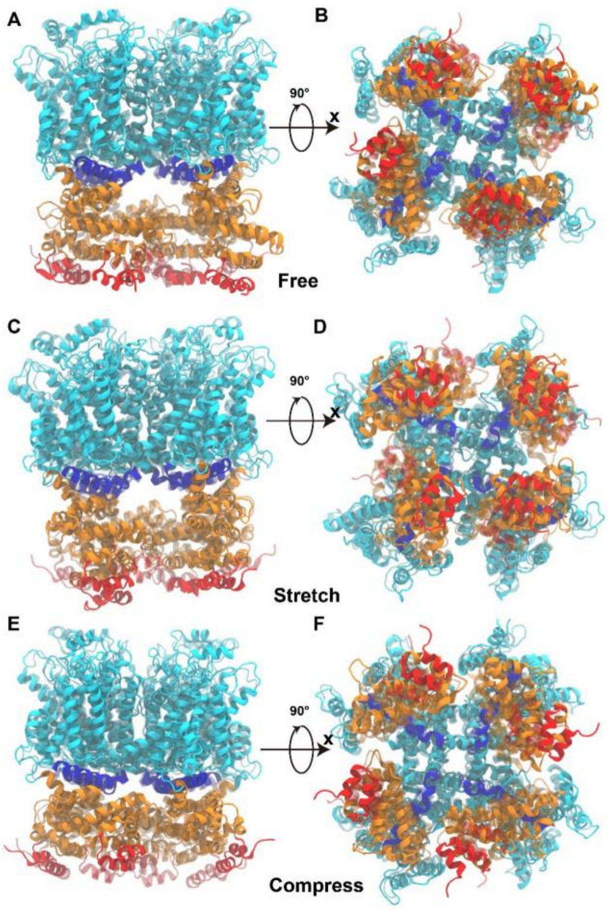


**Figure S6.** The overlaid initial and final structures of the simulation system I in the free, pulling (stretch) and pushing (compress) simulations.

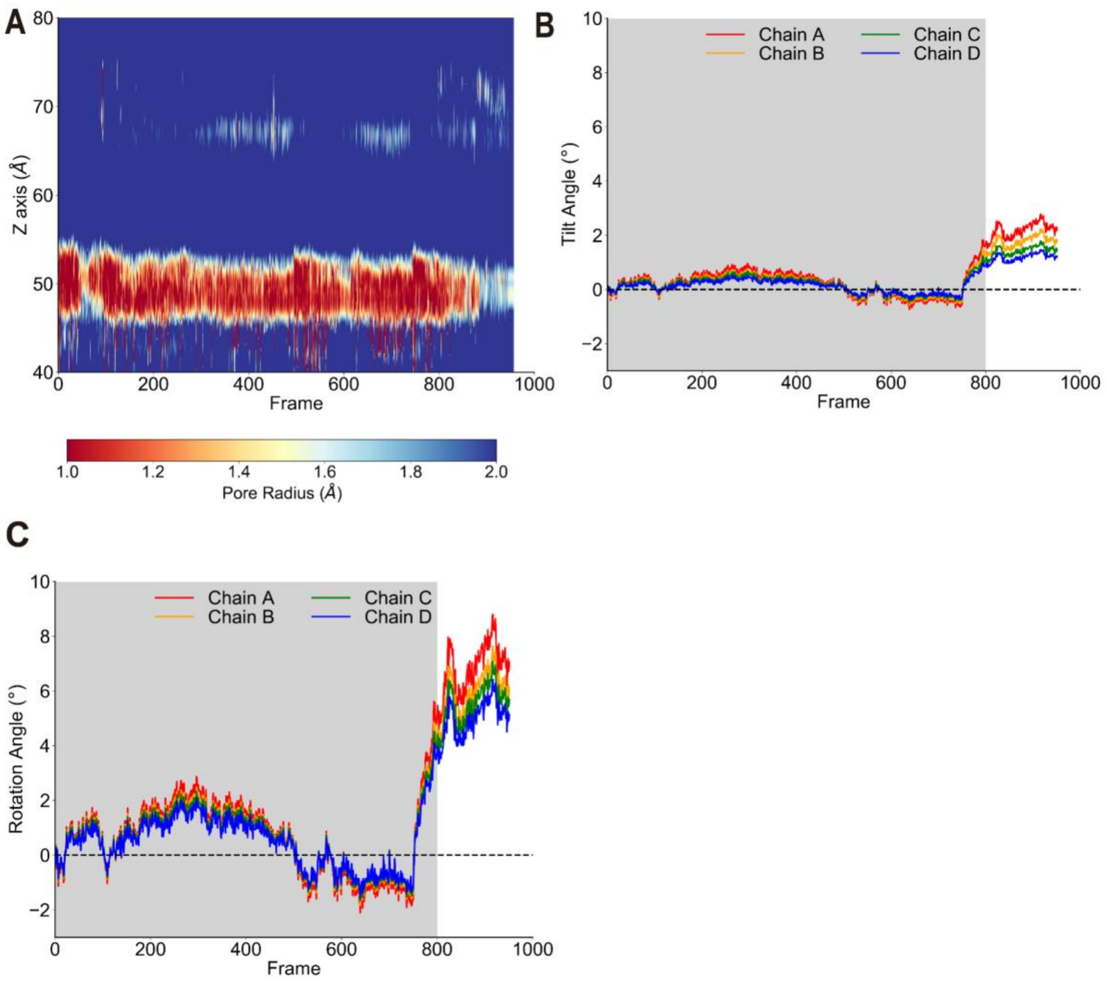


**Figure S7.** The TM pore size evolution and rotation angle evolution of the TRP domain from the principal component analysis (PCA). The trajectories were projected onto the second eigenvector shown in Fig. 2A. **A.** The TM pore size evolution. **B.** The tilt angle evolution of the TRP domain. **C.** The rotation angle evolution of the TRP domain. The lower gate of NompC remains closed in the light gray area.

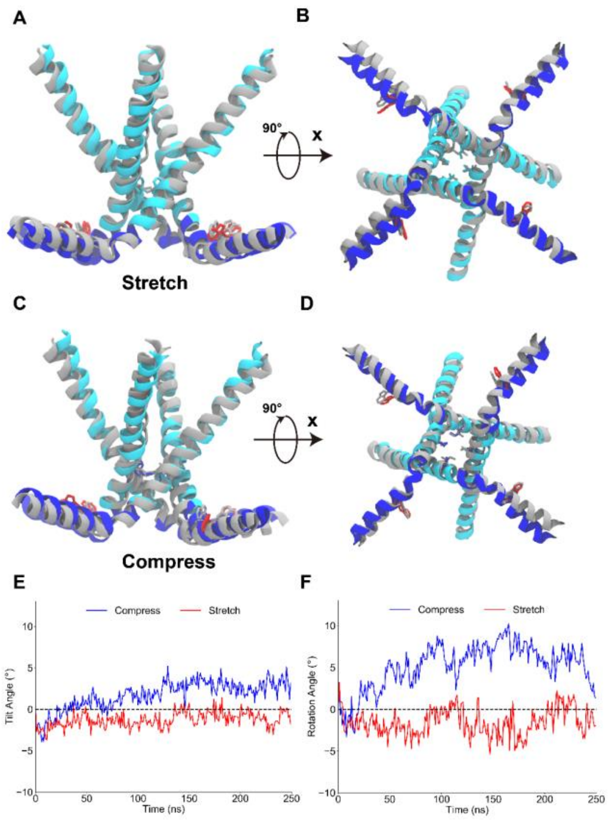


**Figure S8.** The conformational change of the TRP domain in the SMD simulations. The side (**A&C**) and bottom (**B&D**) views of the S6 and TRP domains before (grey) and after (colored) pulling or pushing. The evolution of the average tilt and rotation angles of the TRP domain are shown in **E** and **F** respectively.

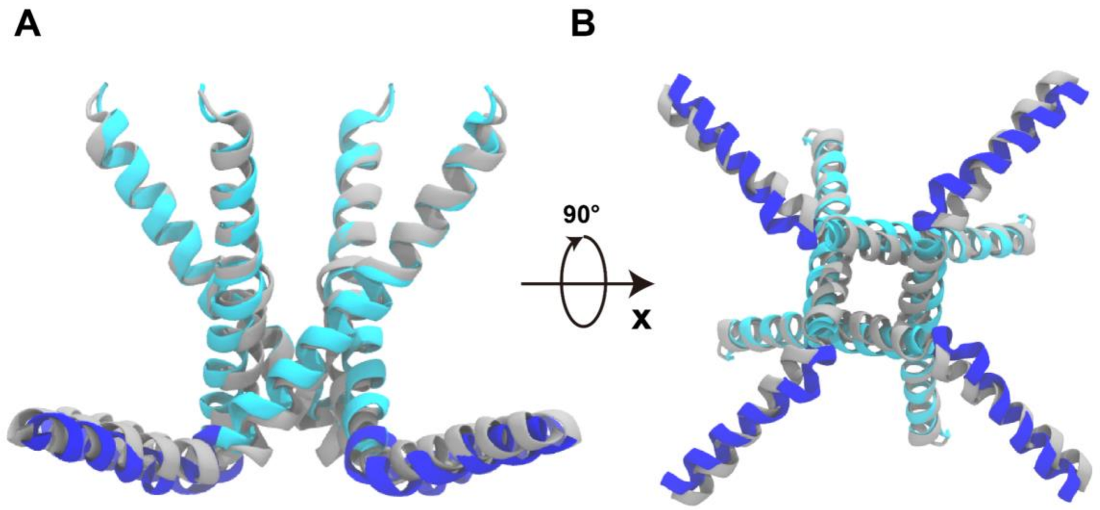


**Figure S9**. The overlaid closed-state (gray) and open-state (blue) structures of TRPV1, obtained in lipid nanodisc (PDB accession numbers: 5irx and 5irz). Only the S6 and TRP domain are shown here for clarity. **A.** From the side view, the TRP domain shows a slight tilt (about 2°) toward the extracellular side when the pore is opened. **B.** From the intracellular view, the TRP domain undergoes a clockwise rotation (about 9°) when the pore is opened.

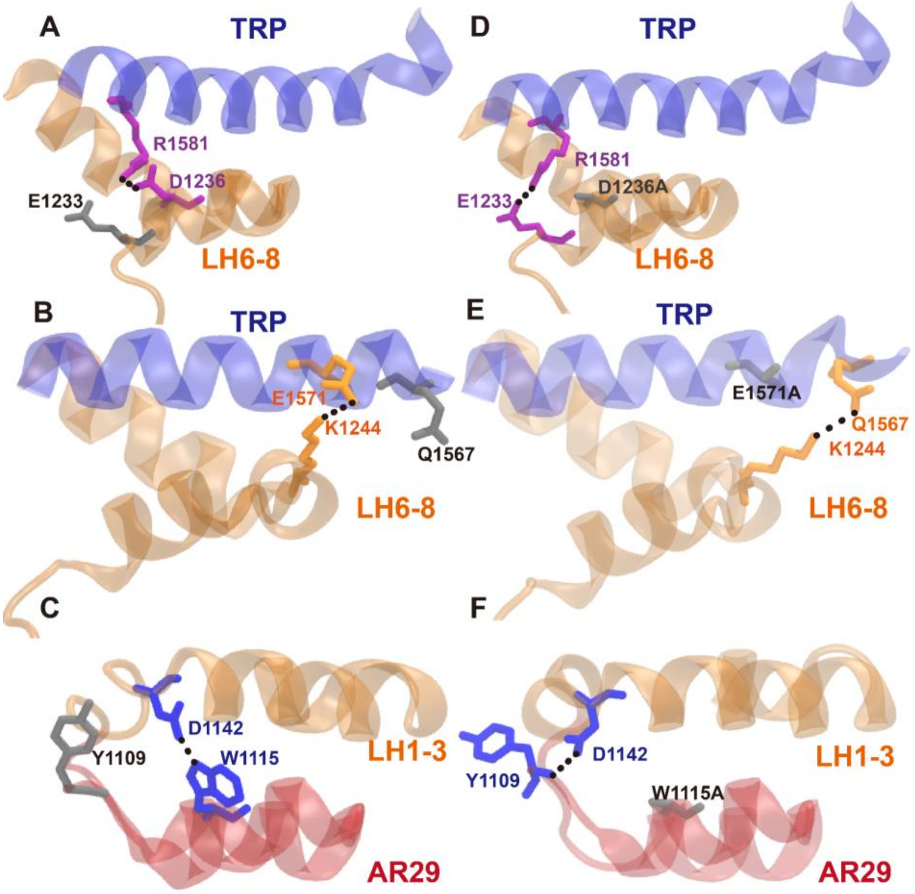


**Figure S10.** The formation of the alternative hydrogen bonds in the mutant, as identified in our MD simulations. Three hydrogen bonds in the wild type NompC around the LH domain (**A, B & C**) can be replaced by alternative hydrogen bonds in the mutant structure (**D, E & F**) to stabilize the interface between the TRP, LH and AR domains, which explains why these mutations did not lead to significant loss-of-function.

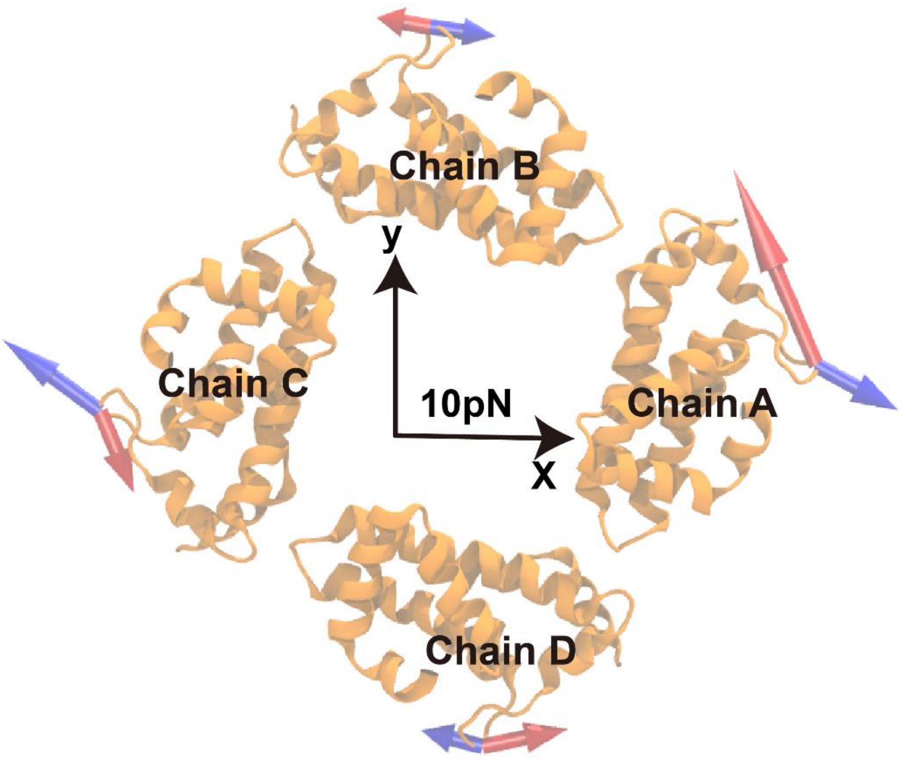


**Figure S11.** The forces on the LH domain projected in the x-y plane. The net forces generated by the stretch (red) and compression (blue) of the AR region were averaged during the 20-40 ns MD/SMD simulations and projected on the x-y plane.

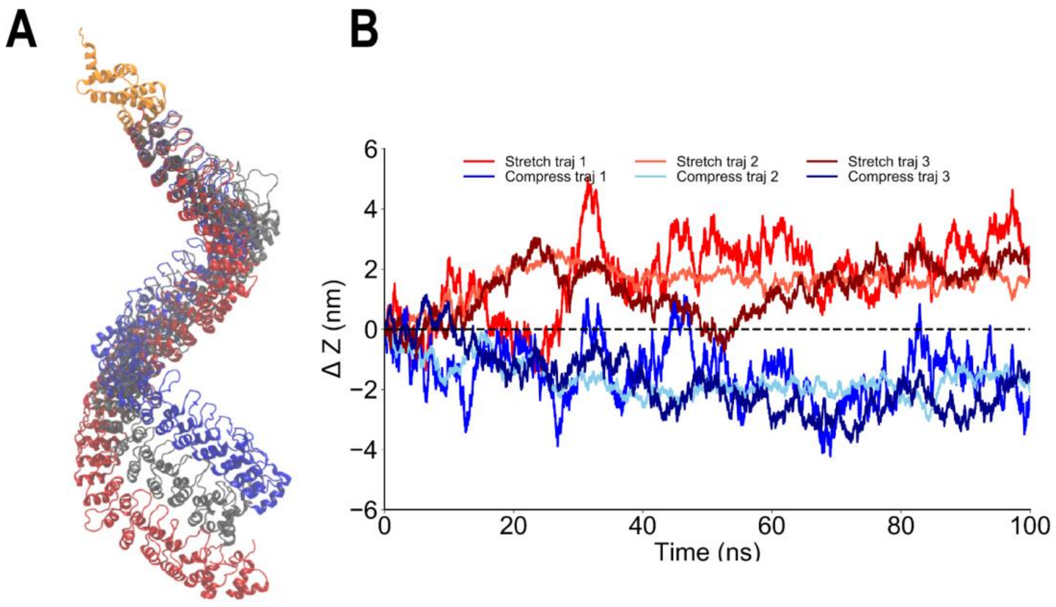


**Figure S12.** Steered MD of the single AR chain of NompC. **A**. The overlaid final structures of the AR chain of NompC in the free (silver) pushing (blue) and pulling (red) simulations. **B.** The change of the AR length from three pulling SMD trajectories (light red, red and dark red) and three pushing SMD trajectories (light blue, blue and dark blue). The change of AR length was measured between the centers of the LH domain and the center of AR1, subtracting the counterpart length from the free MD trajectory.

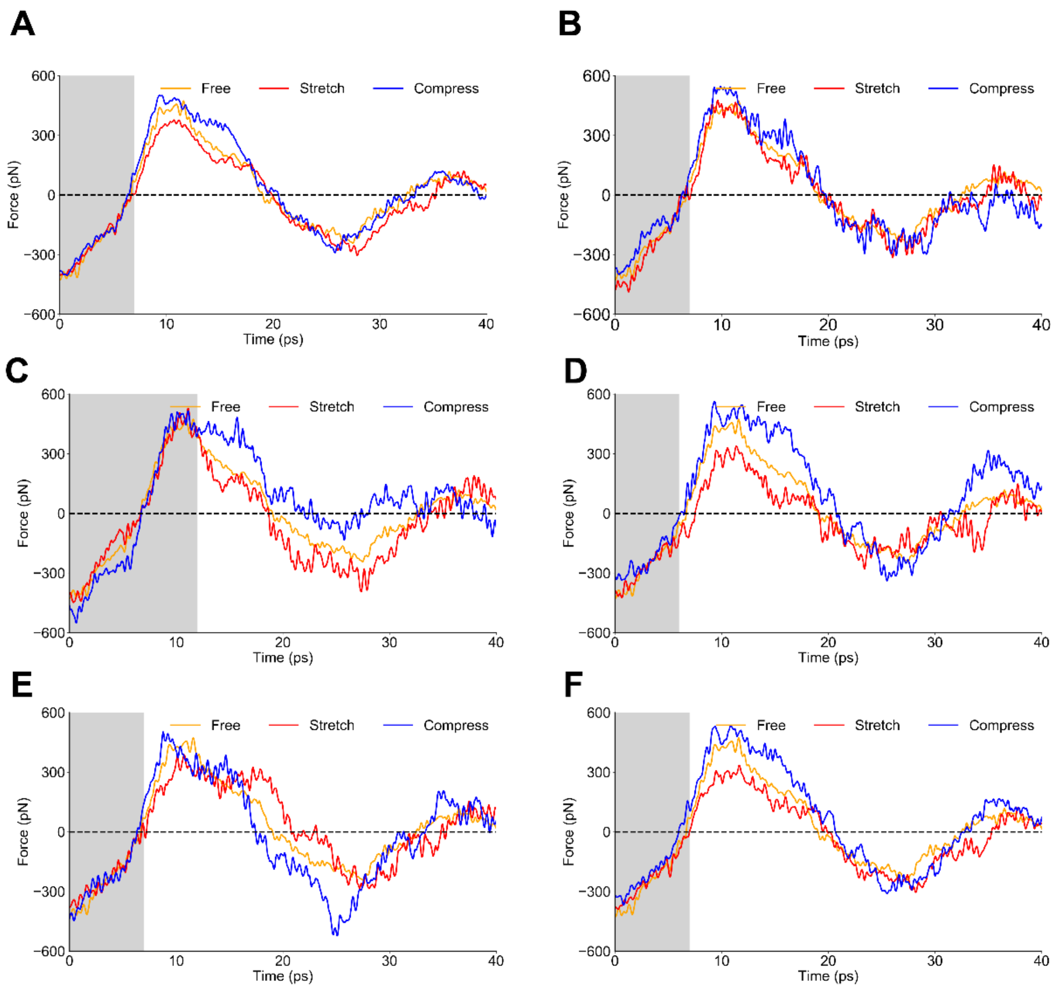


**Figure S13.** The reaction forces to the restraints on the LH domain after applying a force on the AR1**. A**. The averaged force on the LH domain along the z-axis when pulling (red) and pushing (blue) AR1. The grey area indicated the time needed for the force applied on AR1 to impact the LH domain. **B-F,** The force analysis of five independent trajectories estimating the traveling speed of force along the AR region.

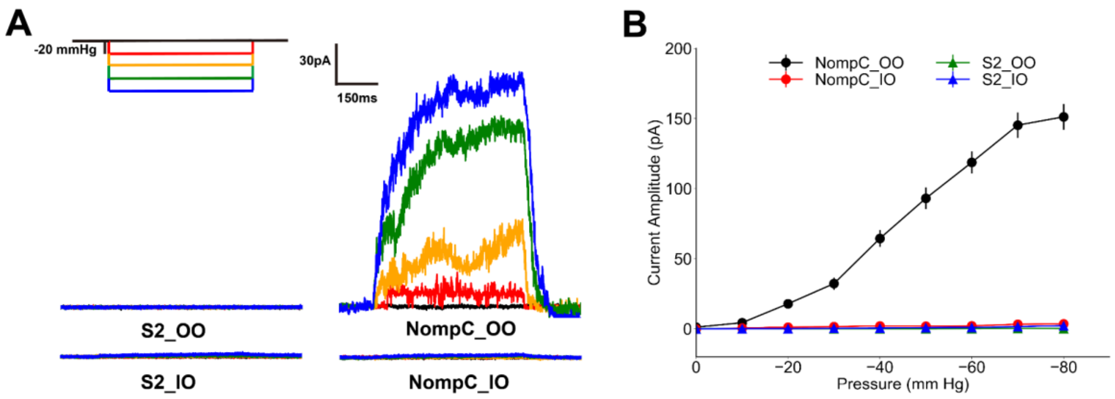


**Figure S14.** Experiment results of the inside-out and outside-out patch-clamp. **A.** Representative traces of mechano-gated current of the S2 Blank Cell and the wild-type NompC under negative pressure ranging from 0 mm Hg to -80 mm Hg, with -20 mm Hg decrements. **B.** Mean mechano-gated current of the wild-type NompC and S2 Blank Cell under negative pressure ranging from 0 mm Hg to -80 mm Hg, with -10 mm Hg decrements.
